# Structuring the unstructured: estimating species-specific absence from multi-species presence data to inform pseudo-absence selection in species distribution models

**DOI:** 10.1101/656629

**Authors:** Simon Croft, Graham C. Smith

**Affiliations:** National Wildlife Management Centre, Animal and Plant Health Agency, Sand Hutton, York, UK

**Keywords:** multi-species, occurrence, opportunistic, pseudo-absence, species distribution modelling, survey effort

## Abstract

1. Species distribution models (SDMs) are an increasingly popular tool in ecology which, together with a vast wealth of data from citizen science projects, have the potential to dramatically improve our understanding of species behaviour for applications such as conservation and wildlife management. However, many of the best performing models require information regarding survey effort, specifically absence, which is typically lacking in opportunistic datasets. To facilitate the use of such models, pseudo-absences from locations without recorded presence must be assumed. Several studies have suggested that survey effort, and hence likely absence, could be estimated from presence-only data by considering records across “target groups” of species defined according to taxonomy.
2. We performed a probabilistic analysis, computing the conditional probability of recording a species given a particular set of species are also recorded, to test the validity of defining target groups by taxonomic order and to explore other potential groupings. Based on this quantification of recording associations we outline a new method to inform pseudo-absence selection comparing predictive performance, measured the area under curve (AUC) statistic, against the standard method of selection across a series of SDMs.
3. Our findings show some support for target grouping classification based on taxonomy but indicate that an alternative classification using survey method may be more appropriate for informing effort and consequently absence. Across 49 terrestrial mammal species, pseudo-absence selection using our proposed method outperformed that of the standard method showing an improvement in the predictive performance of presence-absence models for 17 out of 22 with sufficient data to elicit a significant difference. Based on our method we also observed a substantial improvement in the performance of presence-absence models compared to that of presence-only models (MaxEnt) with a higher AUC for all 22 species showing a significant difference between approaches.
4. We conclude that our method produces sensible robust pseudo-absences which either compliment patterns in known presences or, where conflicts occur, are explainable in terms of ecological variables potentially improving our understanding of species behaviour. Furthermore, we suggest that presence-absence models using these pseudo-absences provide a viable alternative to MaxEnt when modelling using presence-only data.

## Introduction

Knowledge of where species are is important for developing robust wildlife management strategies to address issues of conservation, epidemiology and human-wildlife conflict. In recent years species distribution models (SDMs) have become a popular and potentially powerful tool to inform such investigations (Brotons 2014). These models assess known occurrence against various environmental predictors to establish the ecological niches in which a given species can survive (Elith & Leathwick 2009; Hijmans & Elith 2013). This information can provide valuable insights towards improving our understanding of species ecology which, amongst other outputs, we can use to infer occurrence in areas where no data is available (this includes areas where the species may not currently be present but could exist, which would be important for controlling invasive species). However, many of the best performing models require both positive (presence) and negative (absence) data (Elith *et al.* 2006) which can be difficult to determine reliably without a structured survey; i.e. a known survey effort. Such surveys are typically very expensive especially at a national scale and consequently availability is limited. More commonly, biological data is collected through unstructured opportunistic recording of occurrence. This provides vast amounts of presence data, but absence, or non-detection, is inconsistently reported confounding direct estimation of survey effort and thus true absence.

A common approach to compensate for this deficiency in negative data is to select a random background of pseudo-absence from locations where presence has not been reported. However, it is recognised that this approach is not ideal and that selection should, where possible, be more targeted (Phillips *et al.* 2009; Barbet-Massin *et al.* 2012). Several methods have been suggested which weight selection according to a set of assumptions based on a geographic (Hirzel, Helfer & Metral 2001; Lobo, Jiménez-Valverde & Hortal 2010; Acevedo *et al.* 2012) and/or environmental description (Zaniewski, Lehmann & Overton 2002). More recently it has been argued that aggregating occurrence maps across multiple species could be used to approximate the extent of regular surveillance and therefore absences, i.e. absence of a species can be assumed by non-occurrence when the occurrence of similar species has been recorded (Elith & Leathwick 2007; Phillips *et al.* 2009; Mateo *et al.* 2010; Ranc *et al.* 2016). Similarly, studies using new hierarchical Bayesian SDM frameworks (coupled models describing both habitat suitability and observational detectability; available in R packages such as hSDM (Vieilledent *et al.* 2014) and R-INLA (Rue, Martino & Chopin 2009)) have suggested survey effort (number of visits per site), and thereby non-detection events, could be estimated from presence-only records using the total number of unique visits across associated species, defined in this case as those within the same taxonomic order (Strien, Swaay & Termaat 2013). Whilst both of these approaches offer an intuitive method of inferring negative data based on recorder activity no work has been presented to justify the assumptions regarding species association. It is therefore unclear to what extent an observer recording one species will record another if sighted.

In this study, we aim to explore this problem by analysing recording trends in available opportunistic occurrence data to identify and importantly quantify any links between different species, i.e. given one species is recorded what is the probability of another will also be recorded. From the resulting information we demonstrate a method for pseudo-absence selection based on presence-only data and use it to model the distributions of 49 mammal species across Great Britain. Finally, in the absence of independent structured survey data, as a method of validation for the approach we compare the predictive fit of these models against models generated using an unbiased uniform method of pseudo-absence selection.

## Materials and Methods

### Occurrence data

Occurrence records for all terrestrial mammals between 1995 and 2015 with a minimum resolution of 1000m (coordinate uncertainty <= 1000) were downloaded from the NBN Atlas (http://www.nbnatlas.org) on 05/07/2017 (details of this download are provided in the supporting information; S1). Each record was processed to comprise of a unique “visit”, described by the site (1km OS grid cell) and date (dd/mm/yyyy), and the species observed. Ideally this description of “visit” as a unit of effort would also include a component to distinguish different observers. However, this information is not well represented by the data (less than 10% of records supplied with an observer name; not necessarily unique or well defined) largely due to privacy considerations associated with licensing. In its absence, and to maximise the available data, we assumed that sightings made at the same site on the same day were by the same recorder and consequently duplicate descriptions were removed. Whilst this may not always be accurate we argue that, given the spatial and temporal resolution and the consistency with which the assumption is applied, there is unlikely to be a substantial impact on the outcomes.

### Recording association

To explore recording patterns between species we built an association matrix *A* with each element (*A*_*ij*_) describing the pairwise probability that during any visit species *j* will be recorded (*R*_*j*_) given species *i* is recorded (*R*_*i*_); computed as the number of visits where both were recorded divided by the total number of visits where species *i* was recorded. To remove any bias resulting from differing underlying distributions, visits were only included for sites where both species had been recorded at least once and therefore could be assumed present (for simplicity we assume the probability of false presence to be zero). As such, calculated values in fact reflect the probability of recording species *j* given *i* is recorded and *j* is present (*P*_*j*_); denoted as *p*(*R*_*j*_ | *P*_*j*_ ∩ *R_i_*). We analysed patterns in recording visually using the “heatmap” function in R initially grouping species by taxonomic order and then applying a clustering algorithm to group species by associative similarity. To explore patterns in the recording of species we performed a cluster analysis on this matrix using the “dist” and “hclust” functions in R applying a “manhattan” and “ward.D” method respectively.

### Pseudo-absence selection

If we assume that pairwise associations are independent then, using the probabilities contained in our association matrix, we could compute the probability of recording a species at any site (1km OS grid cell) given it was present and other species had been recorded across a unique set of visits by considering the converse situation, i.e. that a species will not be recorded during any visit despite the recording of other species, as follows:

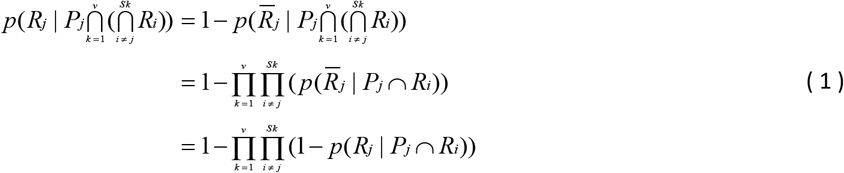

where *v*, *S*_*k*_ and 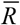 denote the number of visits, the set of species recorded during visit *k* and the non-recording of a species respectively.

However, while this assumption of independence is reasonable between visits it is not sustained between the pairwise interactions of species during a visit (see supporting information for evidence of this dependence; S2). As a consequence we must instead repeat our assessment of recording associations as described previously, but, rather than considering interactions between individual species, we compute the probability of recording a target species given it is present and a given set of species is recorded. In doing so we reduce Equation 1 as follows:

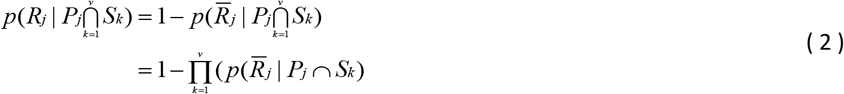

where, as before, *S*_*k*_ denotes the set of species recorded during visit *k*. As the result of these calculations tends to 1 we can be certain that if present a species will be recorded and hence if it has not it must be truly absent. This argument can be expressed mathematically using the laws of conditional probability in the following way:

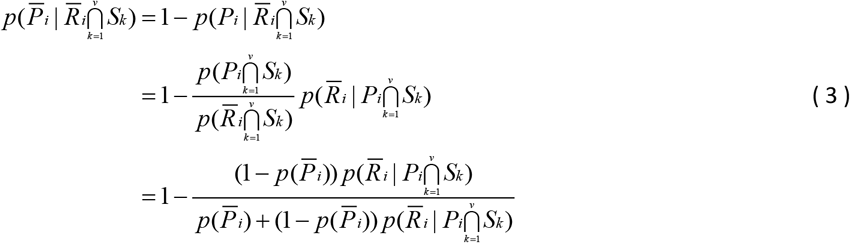

where 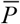 denotes the absence of a species. A full derivation is provided in the supporting information (S3).

### Validation using SDMs

In the absence of an independent presence-absence dataset from a structured survey (with known effort) direct validation of this method is challenging. Nevertheless, we offer some degree of analysis by comparing the impact of pseudo-absence determined as proposed above against a typical uniform random background selection routine on the distribution modelling across a range of species. We argue that an improved area under curve (AUC) statistic indicates some extent of a biologically explainable pattern in these pseudo-absences which reinforces that of the known presence. Whilst this is helpful it should be noted that a reduced AUC is not necessarily evidence that the pseudo-absence selection is invalid, only that some local, spatial or other stochastic effects are not captured in our selection of explanatory variables.

Here, we used an ensemble SDM approach similar to that described by Croft, Chauvenet and Smith (2017). For this approach we first selected our pseudo-absences at random from a background of cells containing at least one occurrence record for any species (an indication that it has been surveyed at least once), excluding cells where our target species is recorded. Each cell in this background was then tested against an assigned probability to determine if it was selected as an absence. Two different probabilities were applied to each cell generating independent datasets: an unbiased selection assuming a uniform probability of true absence equal to 0.02 as suggested by Barbet-Massin *et al.* (2012); and a biased selection based on assumed survey effort across visits with probabilities computed as in Equation 3 assuming the same baseline probability of true absence,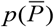.

Once selected we reserved 30% of our presence and absence datasets for testing, accounting for any spatial sorting bias (Hijmans 2012). We used the remaining data to train a selection of SDMs in the R “dismo” package (Hijmans *et al.* 2017), one presence-only model (MaxEnt with default parameters and constrained background selection according to known recording extent) and three presence-absence models (random forest, a support vector machine and a generalised linear model), considering explanatory variables describing climate (worldclim.org; Hijmans *et al.* (2005)), land cover (LCM2007; Morton *et al.* (2011)), topology (altitude; worldclim.org) and human disturbance (population; Office for National Statistics (2015) & National Records of Scotland (2015)). The calibrated AUC for each model was then computed using the reserved testing data and adjusting with respect to the AUC of a random null “geoDist” model (Hijmans 2012). This process was repeated 100 times and the average AUC for each model recorded. We also computed an average of the “best” AUC (highest) for each repetition across the presence-absence models.

Finally, to compare the methods of pseudo-absence selection we considered two main statistics: (i) the difference in average AUC for presence-only MaxEnt models which indicates how well selected absences compliment patterns in the known presence; (ii) the difference in average “best” AUC across the presence-absence models which indicated how informative the absences are in terms of training the model, i.e. explaining the patterns of occurrence to better understand species ecology.

### Results

In order to visualise the relationships described by the computed association matrix we plot values in a heat map where more intense colours reflect a stronger relationship (higher probability). Fig. 1 presents these relationships grouping species by taxonomic order to assess whether there exists a strong justification to adopt this as a target grouping by which to infer recording effort. This appears to suggest that whilst species of the same order are routinely recorded together, particularly bats and deer, there are strong recording associations between species across taxonomic orders.

**Fig. 1:**
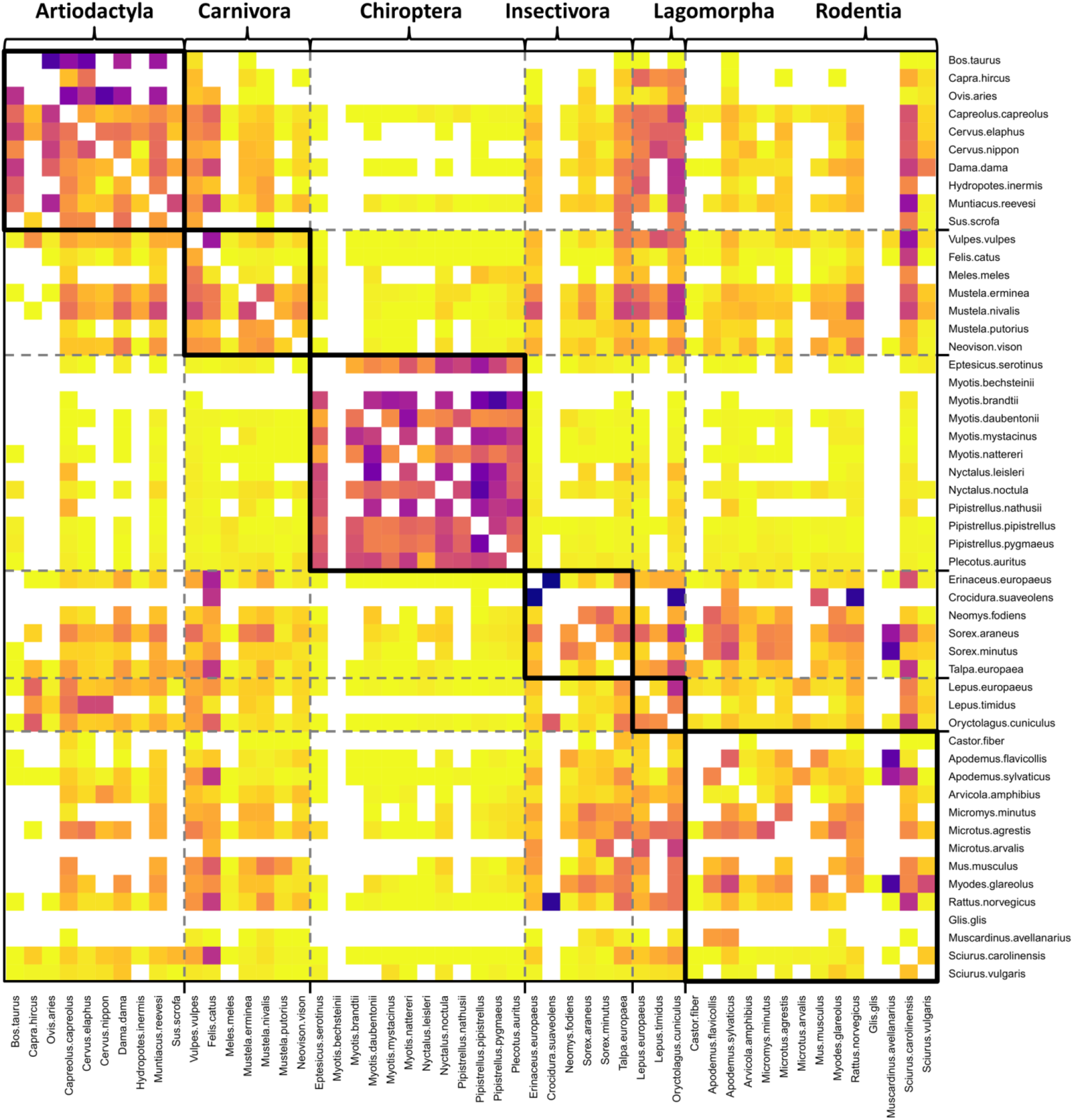
Recording associations by taxonomic order. Heat map visualising a matrix of recording association between species where the value of each cell represents the conditional probability that during any visit (unique 1km OS grid cell and date) species *j* will be recorded given species *i* is recorded (displayed across rows) and both are known to be present. More intense colours denote greater probabilities.

Fig. 2 shows the same associations as in Fig. 1 with species rearranged into groups with similar recording associations as defined by the cluster analysis. Broadly, and perhaps intuitively, these groups can be categorised by the most commonly used survey method. Bats use audio, large mammals use visual and rarer or sensitive species require a specialist continuous monitoring approach such as trapping. For example, common small mammals are often recorded together as many can be surveyed using the same live trapping method.

**Fig. 2:**
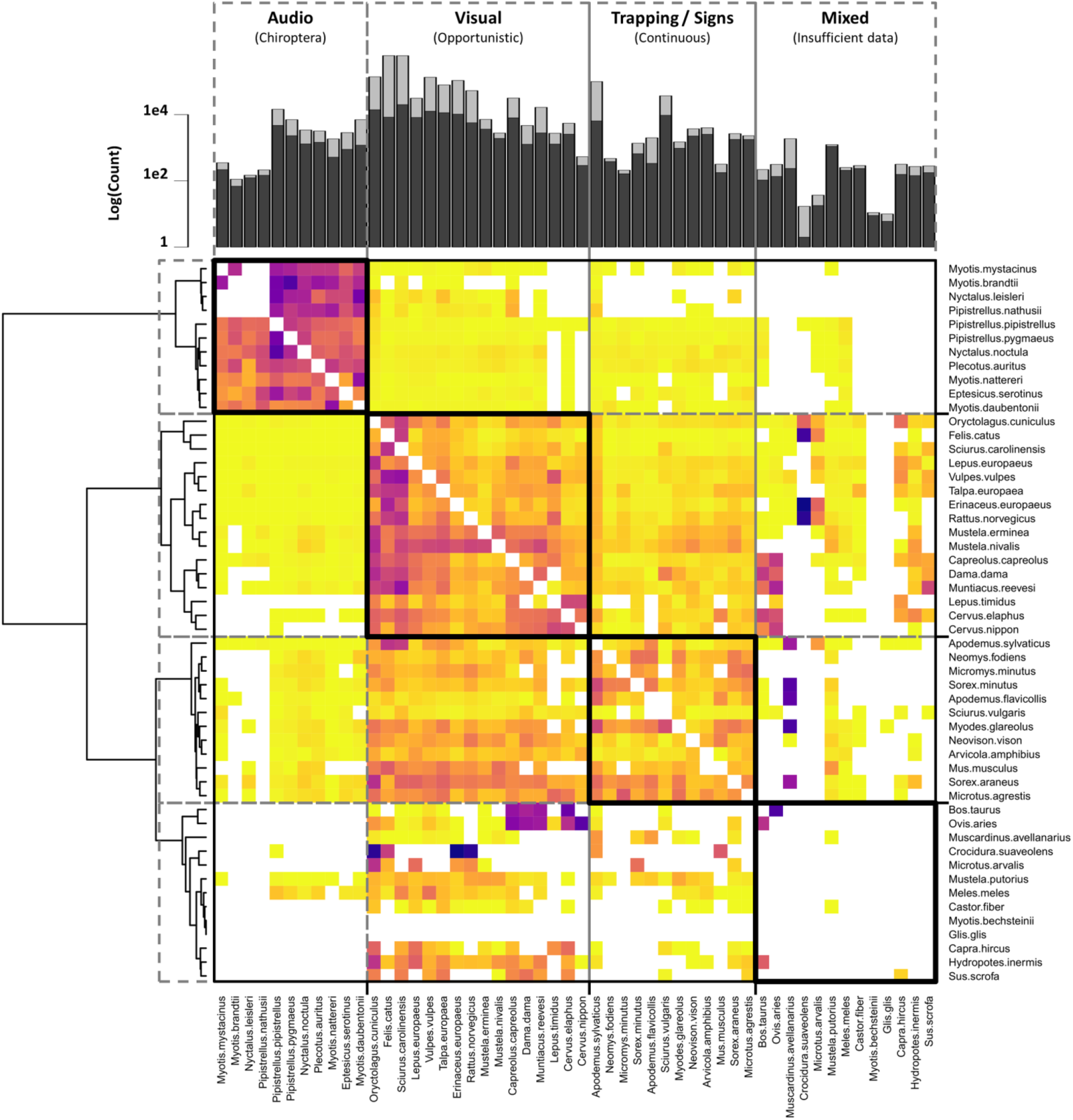
Recording associations by similarity. Heat map visualising a matrix of recording association between species where the value of each cell represents the conditional probability that during any visit (unique 1km OS grid cell and date) species *j* will be recorded given species *i* is recorded (displayed across rows) and both are known to be present. More intense colours denote greater probabilities. Histogram shows the number of records for each species in light grey and the number of unique sites (1km OSGR cells) in dark grey.

Of the 52 species considered 49 had sufficient presence data to perform the species distribution modelling the results of which are outlined in Table 1. We observe that 19 showed a significant difference (measured at the 95% confidence level) in the average AUC for MaxEnt models trained using the unbiased and biased pseudo-absence selection; 12 indicating an improvement in predictive accuracy with the biased selection. For the average “best” AUC across all of the presence-absence methods 22 species showed significant differences; 17 indicating an improvement. Of the separate presence-absence models SVM showed an improvement in 16 out of 20 significant results, random forest 16 out of 21 and GLM 14 out of 21. The majority of significant results were observed for the group of species containing large “common” mammals typically observed visually. All of these results indicated an improvement when using biased pseudo-absence selection with the exception of hedgehogs (*Erinaceus europaeus*) and brown rats (*Rattus norvegicus*); two of the smallest, and potentially most difficult to detect, species in this group.

**Table 1:** Difference in AUC between unbiased and biased pseudo-absence selection. Value indicates improvement in predictive accuracy of model (AUC) based on biased pseudo-absence selection compared with that using an unbiased selection. Asterisks (*) denote the change is significant at the 95% confidence level. Shading highlights groupings by associative similarity.

| Specie | MaxEnt | "Best" PA | SVM | RF | GLM |
| --- | --- | --- | --- | --- | --- |
| *Myotis brandtii* | 0.06 | 0.07 | 0.09 | 0.04 | 0.07 |
| *Myotis mystacinus* | 0.02 | 0.01 | -0.03 | 0.00 | 0.02 |
| *Nyctalus leisleri* | 0.02 | -0.01 | -0.02 | 0.00 | 0.05 |
| *Pipistrellus nathusii* | -0.01 | 0.00 | 0.09 | -0.01 | 0.00 |
| *Nyctalus noctula* | -0.01 | -0.01 | -0.01 | 0.00 | -0.01 |
| *Plecotus auritus* | -0.01 | 0.02 | 0.00 | 0.02 | 0.00 |
| *Pipistrellus pipistrellus* | 0.00 | 0.00 | -0.02 | 0.00 | 0.00 |
| *Pipistrellus pygmaeus* | -0.02* | 0.00 | -0.01 | 0.00 | -0.02 |
| *Myotis nattereri* | 0.03 | 0.04* | -0.06* | 0.05* | 0.04* |
| *Eptesicus serotinus* | 0.00 | 0.01 | -0.01 | 0.01 | 0.01 |
| *Myotis daubentonii* | -0.01 | -0.01 | -0.01 | -0.01 | 0.00 |
| *Bos taurus* | -0.03 | -0.03 | -0.01 | -0.02 | -0.03 |
| *Lepus timidus* | 0.05* | 0.04* | 0.07* | 0.04* | 0.03* |
| *Capra hircus* | 0.02 | 0.05 | 0.06 | 0.06 | 0.06 |
| *Sus scrofa* | 0.09* | 0.13* | 0.00 | 0.13* | 0.11* |
| *Sciurus carolinensis* | -0.02 | 0.01 | 0.04* | 0.01 | -0.03* |
| *Felis catus* | -0.03* | -0.01 | 0.02* | -0.01 | -0.04* |
| *Meles meles* | 0.03 | 0.01 | 0.02 | 0.02 | 0.00 |
| *Mustela putorius* | 0.01 | 0.02 | 0.01 | 0.01 | 0.03 |
| *Arvicola amphibius* | 0.00 | 0.01 | 0.01 | 0.01 | 0.01 |
| *Microtus arvalis* | 0.09 | 0.03 | 0.17 | 0.05 | 0.02 |
| *Sciurus vulgaris* | 0.04* | 0.07* | 0.05* | 0.07* | 0.08* |
| *Muscardinus avellanarius* | 0.00 | -0.06 | -0.12 | -0.04 | -0.06 |
| *Castor fiber* | 0.03 | 0.04 | 0.02 | 0.07 | 0.08 |
| *Myotis bechsteinii* | 0.20 | 0.08 | 0.15 | -0.04 | 0.23 |
| *Cervus nippon* | 0.09* | 0.10* | 0.09* | 0.09* | 0.11* |
| *Hydropotes inermis* | -0.02 | 0.02 | 0.02 | 0.00 | 0.02 |
| *Capreolus capreolus* | 0.09* | 0.13* | 0.16* | 0.13* | 0.14* |
| *Cervus elaphus* | 0.07* | 0.07* | 0.06* | 0.07* | 0.06* |
| *Dama dama* | 0.08* | 0.10* | 0.09* | 0.11* | 0.11* |
| *Muntiacus reevesi* | 0.04* | 0.11* | 0.14* | 0.12* | 0.10* |
| *Erinaceus europaeus* | -0.15* | -0.05* | -0.02* | -0.05* | -0.09* |
| *Rattus norvegicus* | -0.15* | -0.07* | -0.05* | -0.07* | -0.06* |
| *Vulpes vulpes* | -0.14* | 0.06* | 0.08* | 0.07* | -0.02* |
| *Talpa europaea* | 0.06* | 0.17* | 0.21* | 0.17* | 0.17* |
| *Lepus europaeus* | 0.07* | 0.10* | 0.18* | 0.11* | 0.08* |
| *Oryctolagus cuniculus* | 0.00 | 0.12* | 0.19* | 0.13* | 0.11* |
| *Mustela erminea* | 0.05* | 0.06* | 0.08* | 0.06* | 0.07* |
| *Mustela nivalis* | 0.02 | 0.02* | 0.03* | 0.02 | 0.02 |
| *Sorex minutus* | 0.00 | 0.00 | 0.07 | 0.00 | 0.00 |
| *Microtus agrestis* | 0.02* | 0.03* | 0.05* | 0.03* | 0.03* |
| *Neomys fodiens* | -0.04* | -0.05* | -0.02 | -0.05* | -0.02 |
| *Micromys minutus* | 0.01 | 0.02 | 0.05 | 0.04 | 0.03 |
| *Neovison vison* | 0.01 | 0.02 | 0.02 | 0.02 | 0.00 |
| *Apodemus sylvaticus* | -0.14* | -0.12* | -0.07* | -0.11* | -0.12* |
| *Mus musculus* | -0.02 | -0.07* | -0.05 | -0.06* | -0.07* |
| *Apodemus flavicollis* | 0.01 | 0.00 | 0.07 | 0.00 | 0.03 |
| *Sorex araneus* | 0.01 | 0.01 | 0.01 | 0.01 | 0.01 |
| *Myodes glareolus* | 0.02 | 0.03* | 0.02 | 0.03* | 0.03* |

In addition to the comparison in AUCs between absence selection methods we also compared AUC between the presence-only and presence-absence methods within methods. Using the unbiased method of selection our results showed a significant difference between the predictive accuracy of MaxEnt verses the “best” presence-absence for 6 of the 49 species, 5 of which indicated the presence-absence models outperformed MaxEnt. Based on the biased method of absence selection 18 showed a significant difference, all in favour of the presence-absence models over a presence-only MaxEnt model.

## Discussion

The emergence and continued growth of large scale citizen science based repositories for biological records are an important and valuable resource for developing our understanding of species ecology. However, the opportunistic and often unstructured nature of this data means that information regarding survey effort and consequently species absence is lacking. Several recent studies have attempted to address this issue arguing effort can be inferred by considering presence records across sets of species referred to as target groups (Mateo *et al.* 2010; Strien, Swaay & Termaat 2013; Ranc *et al.* 2016). Typically, these groups are defined based on taxonomic descriptions such as order (Strien, Swaay & Termaat 2013). Whilst this may be reasonable for some taxa such as plant and some insects where, due to the nature of observation, recording is typically more comprehensive, the recording of mammal species is generally more ad-hoc and we questioned whether this assumption can be sustained.

In this study we have outlined a probabilistic approach, computing the probability of recording pairs of species during the same visit, to quantify the relationship between recording different species. We use this quantification to assess the validity of defining target groups by taxonomic order and to suggest other possible classifications by which species can be grouped to effectively infer survey effort. Whilst our initial findings presented in Fig. 1 provide some support for a taxonomic grouping, showing high probabilities between species of the same order, we also observed strong relationships between species in different taxonomic orders. When grouping was optimised to cluster species with similar recording associations (Fig. 2) the result, perhaps unsurprisingly, indicate an alternative classification based on common survey method with distinct groups for bats, which are commonly monitored using specialist audio based techniques, large common mammals such as the red fox, which are only really observed through visual observations, and common smaller mammals such as rodents which can be more difficult to detect thus often utilising specialist monitoring such as trapping. For this latter group the species featured can be sub divided into smaller sets which share a common survey methodology explaining why they are often recorded together. There is a further group which contains rare (few records), specialist (species-specific datasets) or sensitive species (poorly recorded at a 1km resolution) for which there is either insufficient data or minimal cross-over to establish relationships with other species. Interestingly, despite their wide distribution and the ease of recording setts, badgers are amongst these species; we suggest that this is largely due to the perception of them as a “sensitive” species so the volume of higher resolution data has been restricted by the recorder.

Whilst the strongest associations between species are concentrated within our identified groups there are some relationships between them suggesting, for example, there is a reasonable probability that where a small mammal is recorded a larger mammal may also be recorded if present. However, the converse appears less likely which is understandable in the context of survey intention as larger mammals can be observed more opportunistically without the need for specialised equipment. A similar argument can be applied to bats which do not show any associations with other species, i.e. do not inform the survey effort for other species, as typically specialist equipment is required and, although large mammals could be observed, most bat recording is conducted during hours of darkness impairing visual observations.

Further analysis of pairwise probabilities showed a dependence on the recording of at least one additional species meaning the probability of recording unique species sets cannot simply be computed as the product of pairwise probabilities. Instead we recomputed conditional probabilities of recording a species given each unique set, multiplying corresponding probabilities across independent visits to bias pseudo-absence selection. In the absence of appropriate data from a structured survey with known effort we proposed a method of validation by comparing the predictive accuracy of models using pseudo-absence based on a standard, unbiased, uniform selection and the biased selection. Looking to the future, we propose that a unique recorder identification across all schemes could be used to improve estimates of effort. Similarly, the increased use of camera trapping, and its integration into presence databases, could include effort and thus determine absence.

In general, where the volume of data was sufficient to produce a significant impact, models trained and evaluated using the biased method of selection showed an improved predictive accuracy in both presence-only and presence-absence models. This suggested that in many cases the absences from this biased method are understandable in terms of the existing species ecology informed by the known presence and simply reinforce those conclusions; hence showing an increase in predictive accuracy for both measures. For a few species we observed a conflict between the presences and selected absences (showing a decrease in predictive accuracy for the presence-only models with testing absence selected according to the biased method when compared with that of the unbiased) but in most cases these conflicts are explained by our chosen variables; improving predictive accuracy of presence-absence models and potentially offering a different perspective in our understanding of the species ecology. There are however some species which indicate a reduced predictive accuracy based on our biased pseudo-absence selection. These tend to be smaller species predominantly recorded near urban areas. This does not necessarily indicate a failure of the method only that likely absences do not fit with patterns of presence and are not explained by our model variables, at least at the spatial scales considered here which may be too large to properly capture the ecological processes of small mammals.

In addition to the comparison between methods of pseudo-absence selection we also compare SDM approaches within each selection, specifically, the performance of presence-only MaxEnt against a presence-absence approach. We would anticipate that if absences are informative then a presence-absence method which makes use of additional information to train the model should have better predictive accuracy. For the unbiased pseudo-absence selection our results show presence-absence models to perform marginally better than MaxEnt in terms of predictive accuracy. However, for our biased selection presence-absence models perform substantially better further validating this method and demonstrating the viability of applying these models to presence-only data rather than simply defaulting to a MaxEnt model.

We accept that refinement may be required especially to account for recording bias which could contribute to these failures (recorder bias has previously been linked to urban areas with higher populations and greater accessibility; Warton, Renner and Ramp (2013)). The method we have proposed could easily be adapted, for example, by applying associations in a different manner to infer study effort for use in hierarchical Bayesian approaches which couple models describing both species ecology and observer behaviour detectability.

The methodology presented here provides a potentially powerful tool to evaluate and understand the patterns in opportunistic recording across all taxa harnessing a vast and ever growing wealth of presence-only data to inform robust analysis of species ecology, particularly with SDMs.

## Supporting information

S1 File (.csv): NBN data providers

S2 File (.pdf): Analysis of dependence between pairwise recording probabilities

S3 File (.pdf): Full derivation of Equation 3

S4 File (.R): Full R script used for analysis

S5 File (.xlsx): Analysis of occurrence records by species

S6 File (.xlsx): Conditional probabilities

S7 File (.xlsx): Breakdown of SDM results

## Acknowledgements

We would like to acknowledge all of the organisations which supplied data to this study via the NBN Atlas. A full list of datasets used in this publication is provided in the supporting information (S1 File).

## Data accessibility

**Occurrence data:** available from NBN Atlas (http://www.nbnatlas.org).

**Environmental data:** (topology & climate) available from worldclim (http://worldclim.org); (land cover) available from CEH (https://eip.ceh.ac.uk) doi: 10.5285/fdf8c8d3-5998-45a5-8431-7f5e6302fc32; (human population) available from Office for National Statistics and National Records of Scotland doi:10.5257/census/aggregate-2011-1.

**R scripts:** uploaded as online supporting information.

**Study output:** uploaded as online supporting information.

## Supporting information

**S1 File (.csv): NBN data providers.** Full list of providers and datasets supplying occurrence records via the NBN Atlas which were used in this study.

**S2 File (.pdf): Analysis of dependence between pairwise recording probabilities.** Heatmap displaying dependence between the pairwise recording of two species and the recording of other species in a particular visit.

**S3 File (.pdf): Full derivation of Equation 3.** Derivation of formula expressing the probability of true absence given all species records.

**S4 File (.R): Full R script used for analysis.** Commented R script detailing methods used to compute association matrix, probability detection maps and perform habitat suitability modelling. Occurrence data and raster maps of explanatory variables are not provided. These can be obtained separately from the sources listed within the main text. Note that file paths will need to be edited in functions if default directory structure is not observed.

**S5 File (.xlsx): Analysis of occurrence records by species.** Excel spreadsheet containing a breakdown of occurrence data by species.

**S6 File (.xlsx): Conditional probabilities.** Excel spreadsheet containing the computed conditional probabilities for both pairwise and set-based associations.

**S7 File (.xlsx): Breakdown of SDM results.** Excel spreadsheet containing a full breakdown by species of the results from the species distribution modelling.

