## Supplementary material for "Structuring the unstructured: estimating species-specific absence from multi-species presence data to inform pseudo-absence selection in species distribution models": S2 File (.pdf): Analysis of dependence between pairwise recording probabilities

**Supporting information file S2: Analysis of dependence between pairwise recording probabilities**

In order to assess the dependance between the recording of pairs of species and the recording of other additional species we performed a chi-squared test on patterns of occurrence as outlined in the R script included as part of the supporting information files (S4). The results of this analysis is shown for each pair of species in the heatmap below. Cells shaded blue denote a chi-squared test with p-value<0.05 suggesting that that recording of the correponding species pair is dependent on at least one other species. Pairwise probabilities computed for cells shaded yellow are considered independent of all other species and hence the probability of a target species being absent where different combination of these associated species are recorded can simply be computed as the product of the pairwise probabilities.

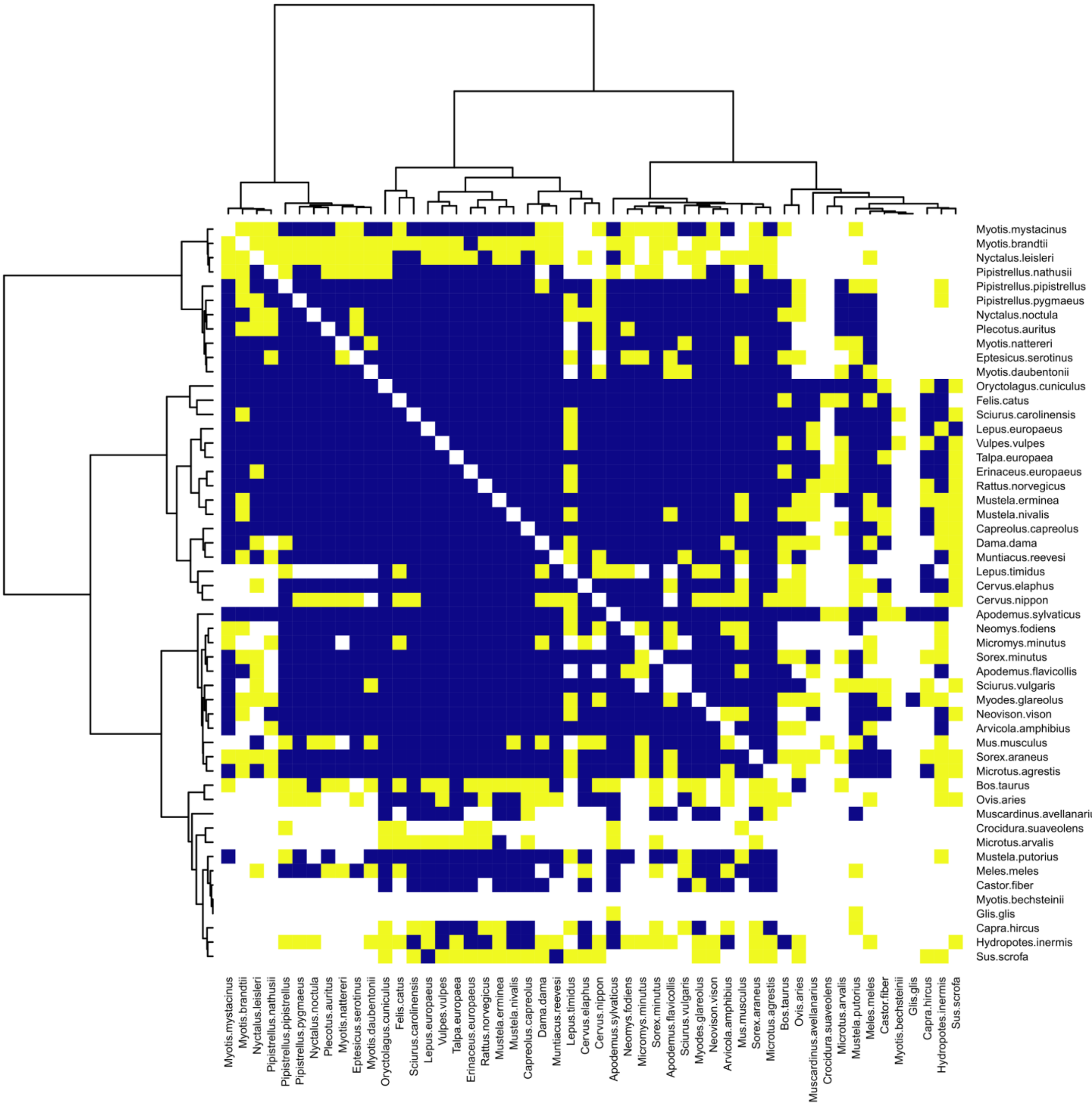


**Fig. S2.1: Dependence between recording species pairs and the recording of other additional species.** Cells coloured blue denote a p-value<0.05 from a chi-squared test of recording patterns suggesting pairwise recording probability is dependent on the recording of at least one other species. Cells coloured yellow indicate pairwise probabilities which are considered independent from the recording of all other species.
