## Supplementary material for "Structuring the unstructured: estimating species-specific absence from multi-species presence data to inform pseudo-absence selection in species distribution models": S3 File (.pdf): Full derivation of Equation 3

**Supporting information file S3: Derivation of the probability of true absence for a species given observation of other species.**

The following formula is a mathematical derivation expressing the probability of true absence $p(\overline{P})$ for a species $i$ given it is not recorded $\overline{R}_i$ but across the time period other species have been recorded with particular frequency $\bigcap_{j\neq i}^{n} R_j$ . This can be written as follows:

$$p(\overline{P}_i \mid \overline{R}_i \bigcap_{j\neq i}^{n} R_j) = 1 - p(P_i \mid \overline{R}_i \bigcap_{j\neq i}^{n} R_j) = 1 - \frac{p(P_i \bigcap \overline{R}_i \bigcap_{j\neq i}^{n} R_j)}{p(\overline{R}_i \bigcap_{j\neq i}^{n} R_j)} = 1 - \frac{p(P_i \bigcap_{j\neq i}^{n} R_j)}{p(\overline{R}_i \bigcap_{j\neq i}^{n} R_j)} p(\overline{R}_i \mid P_i \bigcap_{j\neq i}^{n} R_j) \quad \text{( S3.1 )}$$

If we assume that the probability of a species occurring is independent of other species recording (which we argue is the case when considering species sets across $n$ separate visits) and vice versa, then,

$$p(P_i \bigcap_{j\neq i}^{n} R_j) = p(P_i) p(\bigcap_{j\neq i}^{n} R_j) \quad \text{( S3.2 )}$$

Similarly, if we assume that the probability of not recording a species given it is absence, regardless of other species records is 1, i.e. $p(\overline{R}_i \mid \overline{P}_i \bigcap_{j\neq i}^{n} R_j) = 1$, then we can express the probability of not recording a species given observations of other species as follows:

$$\begin{aligned} p(\overline{R}_i \bigcap_{j\neq i}^{n} R_j) &= p(\overline{P}_i \bigcap \overline{R}_i \bigcap_{j\neq i}^{n} R_j) + p(P_i \bigcap \overline{R}_i \bigcap_{j\neq i}^{n} R_j) \\ &= p(\overline{R}_i \mid \overline{P}_i \bigcap_{j\neq i}^{n} R_j) p(\overline{P}_i \bigcap_{j\neq i}^{n} R_j) + p(\overline{R}_i \mid P_i \bigcap_{j\neq i}^{n} R_j) p(P_i \bigcap_{j\neq i}^{n} R_j) \\ &= p(\overline{P}_i) p(\bigcap_{j\neq i}^{n} R_j) + p(P_i) p(\bigcap_{j\neq i}^{n} R_j) p(\overline{R}_i \mid P_i \bigcap_{j\neq i}^{n} R_j) \end{aligned} \quad \text{( S3.3 )}$$

Therefore we can simplify Eq. S3.1 as follows,

$$\begin{aligned} p(\overline{P}_i \mid \overline{R}_i \bigcap_{j\neq i}^{n} R_j) &= 1 - \frac{p(P_i) p(\bigcap_{j\neq i}^{n} R_j)}{p(\overline{P}_i) p(\bigcap_{j\neq i}^{n} R_j) + p(P_i) p(\bigcap_{j\neq i}^{n} R_j) p(\overline{R}_i \mid P_i \bigcap_{j\neq i}^{n} R_j)} p(\overline{R}_i \mid P_i \bigcap_{j\neq i}^{n} R_j) \\ &= 1 - \frac{p(P_i)}{p(\overline{P}_i) + p(P_i) p(\overline{R}_i \mid P_i \bigcap_{j\neq i}^{n} R_j)} p(\overline{R}_i \mid P_i \bigcap_{j\neq i}^{n} R_j) \end{aligned} \quad \text{( S3.4 )}$$
