## Supplementary material for "Structuring the unstructured: estimating species-specific absence from multi-species presence data to inform pseudo-absence selection in species distribution models": S4 File (.R): Full R script used for analysis: S4 File.R

```
#-------------------------------------------------------------------------------------------------------------------------
# Load libraries
#-------------------------------------------------------------------------------------------------------------------------
library(raster)
library(viridis)
library(usdm)
library(dismo)
library(rJava)
library(kernlab)
library(randomForest)
library(glmulti)
library(boot)

#-------------------------------------------------------------------------------------------------------------------------
# Set environment options
#-------------------------------------------------------------------------------------------------------------------------
continue_on_error<-function(){}
options(error=continue_on_error)
options(stringsAsFactors=F)

#-------------------------------------------------------------------------------------------------------------------------
# Generate and visualise association matrix
#-------------------------------------------------------------------------------------------------------------------------
extent_raster<-raster("./resources/maps/extent/extent.tif") # Import raster of study extent - in our case GB
observations<-read.csv("./resources/data/observations.csv",sep=",",header=T,as.is=T) # Import data - minimum informations (lat,lon,date,species)
observations<-unique(observations[,c("Latitude...processed","Longitude...processed","Event.Date...parsed","Order...matched","Family...matched","Genus...matched","Species...matched")])
colnames(observations)<-c("decimallatitude","decimallongitude","eventdate","order","family","genus","species")
observations[observations==""]<-NA
observations<-observations[complete.cases(observations),] # Clean data - remove incomplete records
coordinates(observations)=~decimallongitude+decimallatitude # Convert to points
proj4string(observations)<-crs("+init=epsg:4326") # Set projection to wgs84
observations<-spTransform(observations,crs(extent_raster)) # Transform coordinate system to match extent raster
observations<-as.data.frame(observations) # Convet back to data frame
observations[,"site"]<-cellFromXY(extent_raster,observations[,c("decimallongitude","decimallatitude")]) # Get unique site location (raster cell) from xy
observations<-observations[as.numeric(format(as.Date(observations[,"eventdate"],format="%Y-%m-%d"),"%Y"))>=1995,] # Data cleaning
observations<-observations[as.numeric(format(as.Date(observations[,"eventdate"],format="%Y-%m-%d"),"%Y"))<=2015,]
observations<-observations[observations[,"order"] %in% c("Artiodactyla","Carnivora","Chiroptera","Insectivora","Lagomorpha","Rodentia"),]
observations<-observations[!(observations[,"family"]=="Phocidae"),]
observations<-observations[!(observations[,"family"]=="Otariidae"),]
observations<-observations[!(observations[,"family"]=="Odobenidae"),]
observations<-observations[!is.na(extract(extent_raster,observations[,"site"])),]
observations<-unique(observations[,c("site","eventdate","order","family","genus","species")])
observations<-observations[order(observations[,"site"],observations[,"eventdate"],observations[,"order"],observations[,"family"],observations[,"genus"],observations[,"species"]),]
observations[,"species"]<-gsub(" ",".",observations[,"species"])
species<-unique(observations[,c("order","family","genus","species")]) # Generate species list
species<-species[order(species[,"order"],species[,"family"],species[,"genus"],species[,"species"]),] # Reorder species by taxonomic description
for(s in 1:nrow(species)){
  specie<-species[s,"species"]
  species[s,"visits"]<-nrow(observations[observations[,"species"]==specie,])
  species[s,"sites"]<-length(unique(observations[observations[,"species"]==specie,"site"]))
} # Compute record summary (records and coverage) for each species
write.csv(species,"../../Results/species.csv",row.names=F)
species<-species[species[,"visits"]>=10,]
observations<-observations[observations[,"species"] %in% species[,"species"],]
# Create matrix of species occurrence per visit
visits<-unique(observations[,c("site","eventdate")])
visits[,"visit"]<-1:nrow(visits)
observations<-merge(observations,visits,by=c("site","eventdate"))
visits[,species[,"species"]]<-0
for(specie in species[,"species"]){visits[observations[observations[,"species"]==specie,"visit"],specie]<-1}
visits<-visits[,-3]
write.csv(visits,"../../Results/visits.csv",row.names=F)
# Create pairwise association matrix
pairwise_matrix<-matrix(NA,nrow=length(species[,"species"]),ncol=length(species[,"species"])) # Initialise matrix
dimnames(pairwise_matrix)<-list(species[,"species"],species[,"species"])
pairwise_matrix<-as.data.frame(pairwise_matrix)
for(specie in species[,"species"]){
  sites<-unique(visits[visits[,specie]==1,"site"])
  data<-visits[visits[,"site"] %in% sites,]
  conditional_species<-species[(species[,"species"]!=specie)&(colSums(data[,species[,"species"]])>=10),"species"]
  for(condition in conditional_species){pairwise_matrix[condition,specie]<-sum(data[,specie]*data[,condition])/sum(data[,condition])}
} # Populate matrix with conditional probabilities
write.csv(pairwise_matrix,"../../Results/pairwise.csv")
pdf("../../Results/pairwise_by_taxonomy.pdf")
  heatmap(t(pairwise_matrix),Rowv=NA,Colv="Rowv",symm=T,col=rev(plasma(100)),cexRow=0.5,cexCol=0.5,margins=c(6,6))
dev.off() # Plot heatmap using taxonomic order (no re-ordering)
nulls<-is.na(pairwise_matrix) # Matrix describing null locations (not enough data to produce estimate)
pairwise_matrix[is.na(pairwise_matrix)]<-0 # Set NAs to 0 
dendro<-hclust(dist(t(pairwise_matrix),method="manhattan"),method="ward.D") # Compute dendrogram to group species by recording similarity
pairwise_matrix[nulls]<-NA Reset NAs
pdf("../../Results/pairwise_by_similarity.pdf")
  heatmap(t(pairwise_matrix),Rowv=as.dendrogram(dendro),Colv="Rowv",symm=T,col=rev(plasma(100)),cexRow=0.5,cexCol=0.5,margins=c(6,6))
dev.off() # Plot heatmap ordering species according to dendrogram
pdf("../../Results/visits_by_similarity.pdf")
  barplot(species[dendro$order,"visits"],names=species[dendro$order,"species"],log="y",ylim=c(1,max(species[,"visits"])),col="grey75")
  barplot(species[dendro$order,"sites"],names=species[dendro$order,"species"],log="y",ylim=c(1,max(species[,"visits"])),col="grey25",add=T)
dev.off() # Plot record count for each species
# Test dependence of pairwise probabilities with the recording of other species
dependence_matrix<-matrix(NA,nrow=length(species[,"species"]),ncol=length(species[,"species"]))
dimnames(dependence_matrix)<-list(species[,"species"],species[,"species"])
dependence_matrix<-as.data.frame(dependence_matrix)
for(specie in species[,"species"]){
  sites<-unique(visits[visits[,specie]==1,"site"])
  data<-visits[visits[,"site"] %in% sites,]
  conditional_species<-species[(species[,"species"]!=specie)&(colSums(data[,species[,"species"]])>0),"species"]
  for(condition in conditional_species){
    table<-table(data[data[,condition]==1,specie],apply(data[data[,condition]==1,species[!(species[,"species"] %in% c(condition,specie)),"species"]],1,paste,collapse=""))
    if(ncol(table)>1){dependence_matrix[condition,specie]<-chisq.test(table)$p.value}
  }
}
write.csv(dependence_matrix,"../../Results/dependence.csv")
nulls<-is.na(dependence_matrix)
dependence_matrix<-1*(dependence_matrix<0.05)
dependence_matrix[nulls]<-NA
pdf("../../Results/dependence_by_similarity.pdf")
  heatmap(t(dependence_matrix),Rowv=as.dendrogram(dendro),Colv="Rowv",symm=T,col=rev(plasma(100)),cexRow=0.5,cexCol=0.5,margins=c(6,6))
dev.off()
# Determine unique conditional species sets
species<-species[,"species"]
recorded_sets<-unique(visits[,species])
sets<-recorded_sets
for(specie in species){
  conditional_sets<-recorded_sets
  conditional_sets[,specie]<-0
  sets<-unique(rbind(sets,conditional_sets[rowSums(conditional_sets)>0,]))
}
# Create set association matrix
sets_matrix<-matrix(NA,nrow=nrow(sets),ncol=length(species))
dimnames(sets_matrix)<-list(apply(sets[,species]==1,1,FUN=function(x)paste(species[x],collapse="_")),species)
sets_matrix<-as.data.frame(sets_matrix)
for(specie in species){
  sites<-unique(visits[visits[,specie]==1,"site"])
  data<-visits[visits[,"site"] %in% sites,]
  conditional_sets<-unique(data[,species])
  conditional_sets[,specie]<-0
  conditional_sets<-unique(conditional_sets[rowSums(conditional_sets)>0,,drop=F])
  if(nrow(conditional_sets)>0){
    species_sets<-conditional_sets
    species_sets[,specie]<-1
    numerators<-table(factor(apply(data[,species],1,paste,collapse=""),levels=apply(species_sets,1,paste,collapse="")))
    denominators<-table(factor(apply(data[,species],1,paste,collapse=""),levels=apply(conditional_sets,1,paste,collapse="")))+numerators
    if(length(denominators[denominators>=10])>0){sets_matrix[apply(conditional_sets[denominators>=10,species]==1,1,FUN=function(x)paste(species[x],collapse="_")),specie]<-numerators[denominators>=10]/denominators[denominators>=10]}
  }
}
write.csv(sets_matrix,"../../Results/sets.csv")

#-------------------------------------------------------------------------------------------------------------------------
# Generate maps of pseudo-absence bias (probability of recording given presence and other species recorded)
#-------------------------------------------------------------------------------------------------------------------------
occurrences<-aggregate(.~site,visits[,-2],sum)
occurrences<-data.frame(xyFromCell(extent_raster,occurrences[,"site"]),occurrences[,-1])
coordinates(occurrences)<-~x+y
occurrences_rasters<-rasterize(occurrences,extent_raster)[[-1]]
occurrences_rasters<-mask(occurrences_rasters,extent_raster)
absence<-visits
absence[,species]<-1
sets_matrix[is.na(sets_matrix)]<-0
for(specie in species){
  conditional_sets<-visits
  conditional_sets[,specie]<-0
  absence[rowSums(conditional_sets[,species])>0,specie]<-1-sets_matrix[apply(conditional_sets[rowSums(conditional_sets[,species])>0,species]==1,1,FUN=function(x)paste(species[x],collapse="_")),specie]
} # For each species compute probabilities of not recording specie given present and "species" recorded
absence<-aggregate(.~site,absence[,-2],function(x) 1-prod(x)) # For each site compute probabilities of recording given all other records (values in each column is 1 minus product of probabilities not recording specie given present and "species" recorded)
absence<-data.frame(xyFromCell(extent_raster,absence[,"site"]),absence[,-1])
coordinates(absence)<-~x+y
absence_rasters<-rasterize(absence,extent_raster)[[-1]]
absence_rasters<-mask(absence_rasters,extent_raster)
trials<-visits
trials[,species]<-0
sets_matrix[is.na(sets_matrix)]<-0
for(specie in species){
  conditional_sets<-visits
  conditional_sets[,specie]<-0
  trials[rowSums(conditional_sets[,species])>0,specie]<-sets_matrix[apply(conditional_sets[rowSums(conditional_sets[,species])>0,species]==1,1,FUN=function(x)paste(species[x],collapse="_")),specie]
}
for(specie in species){trials[,specie]<-pmax(trials[,specie],visits[,specie])}
write.csv(trials,"../../Results/trials.csv",row.names=F)
trials<-aggregate(.~site,trials[,-2],sum)
trials<-data.frame(xyFromCell(extent_raster,trials[,"site"]),trials[,-1])
coordinates(trials)<-~x+y
trials_rasters<-rasterize(trials,extent_raster)[[-1]]
trials_rasters<-mask(trials_rasters,extent_raster)
for(specie in species){
  writeRaster(occurrences_rasters[[which(names(occurrences_rasters)==specie)]],paste("../../Results/occurrence/presence/",specie,".tif",sep=""),format="GTiff",overwrite=T)
  writeRaster(absence_rasters[[which(names(absence_rasters)==specie)]],paste("../../Results/occurrence/absence/",specie,".tif",sep=""),format="GTiff",overwrite=T)
  writeRaster(trials_rasters[[which(names(trials_rasters)==specie)]],paste("../../Results/occurrence/trials/",specie,".tif",sep=""),format="GTiff",overwrite=T)
}

#-------------------------------------------------------------------------------------------------------------------------
# Species distribution modelling
#-------------------------------------------------------------------------------------------------------------------------
predictor_rasters<-stack(paste("./resources/maps/variables/",list.files(path="./resources/maps/variables/",pattern=".tif$"),sep=""))
predictor_data<-as.data.frame(predictor_rasters,xy=T,na.rm=T)
prob_true_absence<-0.02
species<-file_path_sans_ext(list.files(path="../../Results/occurrence/presence",pattern=".tif$"))
for(specie in species){
  presence_raster<-raster(paste("./resources/maps/species/presence/",specie,".tif",sep=""))
  presence_raster[presence_raster==0]<-NA
  presence<-as.data.frame(presence_raster,xy=T,na.rm=T)[,c("x","y")]
  absence_raster<-raster(paste("./resources/maps/species/absence/",specie,".tif",sep=""))
  absence_raster[!is.na(presence_raster)]<-NA
  absence_raster<-1-(((1-prob_true_absence)*(1-absence_raster))/(prob_true_absence+(1-prob_true_absence)*(1-absence_raster))) # Compute map of probability of true absence adjusted for other species records
  absence<-as.data.frame(absence_raster,xy=T,na.rm=T)
  background_raster<-merge(presence_raster,absence_raster)
  # Perform repetions of ensemble method using random uniform pseudo-absence 
  aggregated_suitability_data<-data.frame("count"=rep(0,5),"auc"=rep(NA,5),"auc_sd"=rep(NA,5),"threshold"=rep(NA,5),"threshold_sd"=rep(NA,5),row.names=c("best","maxent","ksvm","rf","glm"))
  aucs<-data.frame("best"=rep(NA,10),"maxent"=rep(NA,10),"ksvm"=rep(NA,10),"rf"=rep(NA,10),"glm"=rep(NA,10))
  thresholds<-rep(NA,10)
  suitability_rasters<-vector(mode="list",length=10)
  presence_rasters<-vector(mode="list",length=10)
  dir.create(paste("../../Results/modelling/uniform/",specie,sep=""))
  dir.create(paste("../../Results/modelling/uniform/",specie,"/runs",sep=""))
  for(i in 1:10){
    dir.create(paste("../../Results/modelling/uniform/",specie,"/runs/",i,sep=""))
    suitability_data<-data.frame("auc"=rep(NA,5),"threshold"=rep(NA,5),row.names=c("null","maxent","ksvm","rf","glm"))
    run_absence<-absence[prob_true_absence>runif(nrow(absence)),c("x","y")] # Select pseudo-absence
    sample_presence<-sample(nrow(presence),floor(0.7*nrow(presence))) # Split presence into training...
    testing_presence<-data.frame(presence[-sample_presence,],"p"=1,extract(predictor_rasters,presence[-sample_presence,])) # ...and testing
    training_presence<-data.frame(presence[sample_presence,],"p"=1,extract(predictor_rasters,presence[sample_presence,]))
    sample_absence<-sample(nrow(run_absence),floor(0.7*nrow(run_absence))) # Split absence into training
    testing_absence<-data.frame(run_absence[-sample_absence,],"p"=0,extract(predictor_rasters,run_absence[-sample_absence,])) # ...and testing
    training_absence<-data.frame(run_absence[sample_absence,],"p"=0,extract(predictor_rasters,run_absence[sample_absence,]))
    pwd<-pwdSample(testing_presence[,c("x","y")],testing_absence[,c("x","y")],training_presence[,c("x","y")],n=1,tr=0.1,lonlat=F,warn=F) # Remove spatial sorting bias in testing data
    testing_presence<-testing_presence[!is.na(pwd[,1]),]
    testing_absence<-testing_absence[na.omit(as.vector(pwd)),]
    testing_presence<-testing_presence[complete.cases(testing_presence),]
    training_presence<-training_presence[complete.cases(training_presence),]
    testing_absence<-testing_absence[complete.cases(testing_absence),]
    training_absence<-training_absence[complete.cases(training_absence),]
    training<-rbind(training_presence,training_absence)
    vif<-vifstep(training[,names(predictor_rasters)],th=10)@results # Remove highly correlated variables
    predictors<-as.vector(vif[order(vif[!is.na(vif[,"VIF"]),"VIF"]),"Variables"])
    if(length(predictors)>31){predictors<-predictors[1:31]}
    suitability_formula<-as.formula(paste("p~",paste(predictors,collapse="+",sep=""),sep=""))
    null<-geoDist(training_presence[,c("x","y")],lonlat=FALSE) # Fit null model to benchmark predictive fit
    eval_null<-evaluate(testing_presence[,c("x","y")],testing_absence[,c("x","y")],null)
    suitability_data["null","auc"]<-eval_null@auc
    xm<-maxent(mask(subset(predictor_rasters,predictors),background_raster),training_presence[,c("x","y")],path=paste("../../output/uniform/",specie,"/",i,"/maxent",sep="")) # Fit maxent model
    eval_xm<-evaluate(testing_presence[,c("x","y")],testing_absence[,c("x","y")],xm,predictor_rasters)
    suitability_data["maxent","auc"]<-eval_xm@auc
    aucs[i,"maxent"]<-suitability_data["maxent","auc"]+0.5-max(0.5,suitability_data["null","auc"])
    svm<-ksvm(suitability_formula,data=training,na.action=na.omit,kpar="automatic") # Fit svm model
    eval_svm<-evaluate(testing_presence,testing_absence,svm)
    suitability_data["ksvm","auc"]<-eval_svm@auc
    aucs[i,"ksvm"]<-suitability_data["ksvm","auc"]+0.5-max(0.5,suitability_data["null","auc"])
    mtry<-tuneRF(training[,predictors],training[,"p"],trace=F,plot=F) # Tune and fit random forest model
    best.m<-mtry[mtry[,2]==min(mtry[, 2]),1]
    rf<-randomForest(suitability_formula,data=training,na.action=na.omit,mtry=best.m)
    eval_rf<-evaluate(testing_presence,testing_absence,rf)
    suitability_data["rf","auc"]<-eval_rf@auc
    aucs[i,"rf"]<-suitability_data["rf","auc"]+0.5-max(0.5,suitability_data["null","auc"])
    glm_multi<-do.call("glmulti",list(suitability_formula,family="binomial",data=training,na.action=na.omit,level=1,method="g",confsetsize=1,deltaB=0,conseq=1,maxit=200,report=F,plotty=F)) # Tune and fit glm
    gl_vector_string<-summary(glm_multi)$bestmodel
    gl_string<-paste(gl_vector_string,sep="",collapse="")
    gl<-glm(gl_string,family="binomial",data=training,na.action=na.omit)
    eval_gl<-evaluate(testing_presence,testing_absence,gl)
    suitability_data["glm","auc"]<-eval_gl@auc
    aucs[i,"glm"]<-suitability_data["glm","auc"]+0.5-max(0.5,suitability_data["null","auc"])
    best<-rownames(suitability_data)[order(-suitability_data[3:5,"auc"])[1]+2] # Identify best predictive model based on auc
    # Predict suitaility raster across full variable extent
    switch(best,
      "maxent"={
        suitability_raster<-predict(xm,predictor_rasters)
        suitability_data["maxent","threshold"]<-eval_xm@t[which.max(eval_xm@TPR+eval_xm@TNR)]
      },
      "ksvm"={
        suitability_raster<-rasterFromXYZ(cbind(predictor_data[,c("x","y")],predict(svm,predictor_data)))
        suitability_data["ksvm","threshold"]<-eval_svm@t[which.max(eval_svm@TPR+eval_svm@TNR)]
      },
      "rf"={
        suitability_raster<-rasterFromXYZ(cbind(predictor_data[,c("x","y")],predict(rf,predictor_data)))
        suitability_data["rf","threshold"]<-eval_rf@t[which.max(eval_rf@TPR+eval_rf@TNR)]
      },
      "glm"={
        suitability_raster<-rasterFromXYZ(cbind(predictor_data[,c("x","y")],predict(gl,predictor_data,type="response")))
        suitability_data["glm","threshold"]<-inv.logit(eval_gl@t[which.max(eval_gl@TPR+eval_gl@TNR)])
      }
    )
    aggregated_suitability_data["best","count"]<-aggregated_suitability_data["best","count"]+1
    aggregated_suitability_data[best,"count"]<-aggregated_suitability_data[best,"count"]+1
    aucs[i,"best"]<-aucs[i,best]
    # Standardise raster and threshold
    min<-minValue(suitability_raster)
    max<-maxValue(suitability_raster)
    suitability_data[best,"threshold"]<-(suitability_data[best,"threshold"]-min)/(max-min)
    thresholds[i]<-suitability_data[best,"threshold"]
    write.csv(suitability_data,file=paste("../../Results/modelling/uniform/",specie,"/runs/",i,"/suitability.csv",sep=""))
    suitability_raster<-(suitability_raster-min)/(max-min)
    suitability_raster<-extend(suitability_raster,extent(predictor_rasters[[1]]))
    writeRaster(suitability_raster,paste("../../Results/modelling/uniform/",specie,"/runs/",i,"/suitability.tif",sep=""),format="GTiff",overwrite=T)
    suitability_rasters[[i]]<-suitability_raster
    presence_rasters[[i]]<-suitability_raster>=thresholds[i] # Convert suitability raster to binary output
  }
  # Aggregate repetition output into single output
  for(row in rownames(aggregated_suitability_data)){
    aggregated_suitability_data[row,"auc"]<-mean(aucs[,row])
    aggregated_suitability_data[row,"auc_sd"]<-sd(aucs[,row])
  }
  aggregated_suitability_data["best","threshold"]<-mean(thresholds)
  aggregated_suitability_data["best","threshold_sd"]<-sd(thresholds)
  write.csv(aggregated_suitability_data,file=paste("../../Results/modelling/uniform/",specie,"/suitability.csv",sep=""))
  suitability_stack<-stack(suitability_rasters)
  writeRaster(calc(suitability_stack,mean),filename=paste("../../Results/modelling/uniform/",specie,"/suitability.tif",sep=""),format="GTiff",overwrite=T)
  writeRaster(calc(suitability_stack,sd),filename=paste("../../Results/modelling/uniform/",specie,"/uncertainty.tif",sep=""),format="GTiff",overwrite=T)
  presence_stack<-stack(presence_rasters)
  writeRaster(calc(presence_stack,mean),filename=paste("../../Results/modelling/uniform/",specie,"/presence.tif",sep=""),format="GTiff",overwrite=T)
  # Perform repetitions of ensemble method using random biased pseudo-absence
  aggregated_suitability_data<-data.frame("count"=rep(0,5),"auc"=rep(NA,5),"auc_sd"=rep(NA,5),"threshold"=rep(NA,5),"threshold_sd"=rep(NA,5),row.names=c("best","maxent","ksvm","rf","glm"))
  aucs<-data.frame("best"=rep(NA,10),"maxent"=rep(NA,10),"ksvm"=rep(NA,10),"rf"=rep(NA,10),"glm"=rep(NA,10))
  thresholds<-rep(NA,10)
  suitability_rasters<-vector(mode="list",length=10)
  presence_rasters<-vector(mode="list",length=10)
  dir.create(paste("../../Results/modelling/weighted/",specie,sep=""))
  dir.create(paste("../../Results/modelling/weighted/",specie,"/runs",sep=""))
  for(i in 1:10){
    suitability_data<-data.frame("auc"=rep(NA,5),"threshold"=rep(NA,5),row.names=c("null","maxent","ksvm","rf","glm"))
    run_absence<-absence[absence[,3]>runif(nrow(absence)),c("x","y")] # Select pseudo-absence by testing each cell value (probability of true absence) against random value from uniform distribution on [0,1]
    sample_presence<-sample(nrow(presence),floor(0.7*nrow(presence)))
    testing_presence<-data.frame(presence[-sample_presence,],"p"=1,extract(predictor_rasters,presence[-sample_presence,]))
    training_presence<-data.frame(presence[sample_presence,],"p"=1,extract(predictor_rasters,presence[sample_presence,]))
    sample_absence<-sample(nrow(run_absence),floor(0.7*nrow(run_absence)))
    testing_absence<-data.frame(run_absence[-sample_absence,],"p"=0,extract(predictor_rasters,run_absence[-sample_absence,]))
    training_absence<-data.frame(run_absence[sample_absence,],"p"=0,extract(predictor_rasters,run_absence[sample_absence,]))
    pwd<-pwdSample(testing_presence[,c("x","y")],testing_absence[,c("x","y")],training_presence[,c("x","y")],n=1,tr=0.1,lonlat=F,warn=F)    
    testing_presence<-testing_presence[!is.na(pwd[,1]),]
    testing_absence<-testing_absence[na.omit(as.vector(pwd)),]
    testing_presence<-testing_presence[complete.cases(testing_presence),]
    training_presence<-training_presence[complete.cases(training_presence),]
    testing_absence<-testing_absence[complete.cases(testing_absence),]
    training_absence<-training_absence[complete.cases(training_absence),]
    training<-rbind(training_presence,training_absence)
    vif<-vifstep(training[,names(predictor_rasters)],th=10)@results
    predictors<-as.vector(vif[order(vif[!is.na(vif[,"VIF"]),"VIF"]),"Variables"])
    if(length(predictors)>31){predictors<-predictors[1:31]}
    suitability_formula<-as.formula(paste("p~",paste(predictors,collapse="+",sep=""),sep=""))
    null<-geoDist(training_presence[,c("x","y")],lonlat=FALSE)
    eval_null<-evaluate(testing_presence[,c("x","y")],testing_absence[,c("x","y")],null)
    suitability_data["null","auc"]<-eval_null@auc
    xm<-maxent(mask(subset(predictor_rasters,predictors),background_raster),training_presence[,c("x","y")],path=paste("../../output/weighted/",specie,"/",i,"/maxent",sep=""))
    eval_xm<-evaluate(testing_presence[,c("x","y")],testing_absence[,c("x","y")],xm,predictor_rasters)
    suitability_data["maxent","auc"]<-eval_xm@auc
    aucs[i,"maxent"]<-suitability_data["maxent","auc"]+0.5-max(0.5,suitability_data["null","auc"])
    svm<-ksvm(suitability_formula,data=training,na.action=na.omit,kpar="automatic")
    eval_svm<-evaluate(testing_presence,testing_absence,svm)
    suitability_data["ksvm","auc"]<-eval_svm@auc
    aucs[i,"ksvm"]<-suitability_data["ksvm","auc"]+0.5-max(0.5,suitability_data["null","auc"])
    mtry<-tuneRF(training[,predictors],training[,"p"],trace=F,plot=F)
    best.m<-mtry[mtry[,2]==min(mtry[, 2]),1]
    rf<-randomForest(suitability_formula,data=training,na.action=na.omit,mtry=best.m)
    eval_rf<-evaluate(testing_presence,testing_absence,rf)
    suitability_data["rf","auc"]<-eval_rf@auc
    aucs[i,"rf"]<-suitability_data["rf","auc"]+0.5-max(0.5,suitability_data["null","auc"])
    glm_multi<-do.call("glmulti",list(suitability_formula,family="binomial",data=training,na.action=na.omit,level=1,method="g",confsetsize=1,deltaB=0,conseq=1,maxit=200,report=F,plotty=F))
    gl_vector_string<-summary(glm_multi)$bestmodel
    gl_string<-paste(gl_vector_string,sep="",collapse="")
    gl<-glm(gl_string,family="binomial",data=training,na.action=na.omit)
    eval_gl<-evaluate(testing_presence,testing_absence,gl)
    suitability_data["glm","auc"]<-eval_gl@auc
    aucs[i,"glm"]<-suitability_data["glm","auc"]+0.5-max(0.5,suitability_data["null","auc"])
    best<-rownames(suitability_data)[order(-suitability_data[3:5,"auc"])[1]+2]
    switch(best,
      "maxent"={
        suitability_raster<-predict(xm,predictor_rasters)
        suitability_data["maxent","threshold"]<-eval_xm@t[which.max(eval_xm@TPR+eval_xm@TNR)]	
      },
      "ksvm"={
        suitability_raster<-rasterFromXYZ(cbind(predictor_data[,c("x","y")],predict(svm,predictor_data)))
        suitability_data["ksvm","threshold"]<-eval_svm@t[which.max(eval_svm@TPR+eval_svm@TNR)]	
      },
      "rf"={	
        suitability_raster<-rasterFromXYZ(cbind(predictor_data[,c("x","y")],predict(rf,predictor_data)))
        suitability_data["rf","threshold"]<-eval_rf@t[which.max(eval_rf@TPR+eval_rf@TNR)]
      },
      "glm"={
        suitability_raster<-rasterFromXYZ(cbind(predictor_data[,c("x","y")],predict(gl,predictor_data,type="response")))
        suitability_data["glm","threshold"]<-inv.logit(eval_gl@t[which.max(eval_gl@TPR+eval_gl@TNR)])
      }
    )
    aggregated_suitability_data["best","count"]<-aggregated_suitability_data["best","count"]+1
    aggregated_suitability_data[best,"count"]<-aggregated_suitability_data[best,"count"]+1
    aucs[i,"best"]<-aucs[i,best]
    min<-minValue(suitability_raster)
    max<-maxValue(suitability_raster)
    suitability_data[best,"threshold"]<-(suitability_data[best,"threshold"]-min)/(max-min)
    thresholds[i]<-suitability_data[best,"threshold"]
    write.csv(suitability_data,file=paste("../../Results/modelling/weighted/",specie,"/runs/",i,"/suitability.csv",sep=""))
    suitability_raster<-(suitability_raster-min)/(max-min)
    suitability_raster<-extend(suitability_raster,extent(predictor_rasters[[1]]))
    writeRaster(suitability_raster,paste("../../Results/modelling/weighted/",specie,"/runs/",i,"/suitability.tif",sep=""),format="GTiff",overwrite=T)
    suitability_rasters[[i]]<-suitability_raster  
    presence_rasters[[i]]<-suitability_raster>=thresholds[i]
  }
  for(row in rownames(aggregated_suitability_data)){
    aggregated_suitability_data[row,"auc"]<-mean(aucs[,row])
    aggregated_suitability_data[row,"auc_sd"]<-sd(aucs[,row])
  }
  aggregated_suitability_data["best","threshold"]<-mean(thresholds)
  aggregated_suitability_data["best","threshold_sd"]<-sd(thresholds)
  write.csv(aggregated_suitability_data,file=paste("../../Results/modelling/weighted/",specie,"/suitability.csv",sep=""))
  suitability_stack<-stack(suitability_rasters)
  writeRaster(calc(suitability_stack,mean),filename=paste("../../Results/modelling/weighted/",specie,"/suitability.tif",sep=""),format="GTiff",overwrite=T)
  writeRaster(calc(suitability_stack,sd),filename=paste("../../Results/modelling/weighted/",specie,"/uncertainty.tif",sep=""),format="GTiff",overwrite=T)
  presence_stack<-stack(presence_rasters)
  writeRaster(calc(presence_stack,mean),filename=paste("../../Results/modelling/weighted/",specie,"/presence.tif",sep=""),format="GTiff",overwrite=T)
}
```
